# Cell-based calcification assays – magnesium overrules calcium phosphate saturation in overall outcome

**DOI:** 10.64898/2025.12.23.696208

**Authors:** Aaron Morgan, Christian Hasberg, Camilla Winkler, Robert Dzhanaev, Steffen Gräber, Luca Reicher, Andrea Gorgels, Claudia Goettsch, Willi Jahnen-Dechent

## Abstract

Soft tissue calcification, a common cause of cardiovascular mortality in conditions like chronic kidney disease (CKD), is influenced by magnesium (Mg). However, the exact mechanism, whether extracellular chemistry-mediated or cell-mediated, remains unclear. Here we focused on the extracellular milieu. Using fluorescence-labelled fetuin-A and a live imaging platform, we found that Mg reduced spontaneous mineral precipitation by stabilizing calcium and phosphate as colloidal protein-mineral particles (CPP) in calcification media containing up to 10 mM calcium and phosphate. Addition of 1.25-5 mM Mg progressively stabilized the protein-mineral particles in the fluid phase, increasing calcification. Dynamic light scattering revealed smaller and more numerous CPP at up to 4 mM Mg. At 10 mM Mg, no calcification occurred even with 10 mM added calcium and phosphate. This suggested a critical Mg concentration range in supersaturated calcification media, where Mg paradoxically enhanced calcification by stabilizing mineral precursors and increasing their availability. These findings demonstrate Mg’s dual role: an inhibitor of protein-mineral complex aggregation at high concentrations and a facilitator of mineral availability from colloidal protein-mineral phases at concentrations up to about 5 mM Mg. This highlights the role of Mg in providing highly dynamic, mineral-rich, stable protein-mineral complexes driving matrix calcification, yet preventing mineral precipitation.

## Introduction

Soft tissue calcification, a pathology facilitated by dysregulated mineral metabolism in chronic kidney disease (CKD), correlates strongly with cardiovascular morbidity and mortality [1, 2]. Calcium and phosphate drive calcification while, in many respects, magnesium counteracts the ill effects of calcium [3]. Clinical studies associate hypomagnesemia with increased coronary artery calcification [4, 5], cardiovascular mortality, and sudden cardiac death [6, 7], yet mechanistic insights into the role of Mg in calcification remain sparse.

Magnesium reduces the crystallinity of calcium phosphate and stabilizes amorphous calcium phosphate [8], delaying hydroxyapatite crystallization [9]. Magnesium generally acts to promote cell survival by stabilizing cell membranes, reducing oxidative stress, and inhibiting excessive calcium influx into cells, and thus apoptosis [3]. Cell culture studies suggest that Mg suppresses phosphate-induced osteogenic differentiation of vascular smooth muscle cells (VSMC) [10]. Additionally, in VSMC, Mg inhibits osteogenic signaling mediated by Wnt/β-catenin [11] and Osterix [12]. Recent research suggests that protein-mineral complexes called calciprotein particles (CPP) regulated in vitro collagen fiber mineralization. In particular, early unstable complexes called calciprotein monomers (CPM) or transient CPP (tCPP) mineralized collagen fibers much better than more stable secondary calciprotein particles (CPP-2) [13]. Mg is known to stabilize amorphous CPP-1 thereby delaying their transition into crystalline CPP-2 [14]. Whether Mg also stabilizes CPM and tCPP is unknown.

Ionic and topological shielding are two mechanisms known to stabilize protein-mineral complexes like CPP. Ionic shielding is predominantly the adsorption of Mg to the surface of nascent mineral clusters, which delays the phase transition to crystalline hydroxyapatite [15]. Serum-derived mineral chaperone proteins like fetuin-A [16] form a protein shell that sterically encases the mineral core providing topological shielding [17]. This physical barrier prevents the spontaneous propagation of the mineral and the maturation of CPP, thereby contributing to the colloidal stability of the suspension. While both mechanisms effectively inhibit spontaneous precipitation, they increase the lifetime and availability of mineral precursors, potentially enhancing their delivery to calcifiable matrix.

Fluorescence-labelled fetuin-A has enabled live calcification imaging in cell culture [18] and real-time tracking of CPP in live mice [19]. In this work, live imaging was combined with vascular smooth muscle cell (VSMC) culture models to investigate the role of Mg in stabilizing calcification fluids and in cell-mediated calcification. We hypothesized that Mg delays nucleation and particle aggregation thus stabilizing amorphous calcium phosphate and immature CPP enabling sustained ECM-mineral interactions. We show that this was indeed the case and we present a rationale for optimizing calcification fluids for the study of cell, extracellular matrix, and biomaterial scaffold calcification.

## Methods and Materials

### T50 calcification propensity test

The calcification propensity of serum protein-containing solutions was determined by measuring the T50 time, a functional metric defined as the time required for a supersaturated solution of Ca and Pi to reach 50% of a sudden increase in turbidity, signifying the transformation of primary CPP/CPP-1 into secondary CPP/CPP-2 [14]. Four pre-warmed (34.5°C) stock solutions were prepared: a 140 mM NaCl saline solution, a Ca stock solution (40 mM CaCl₂, 100 mM HEPES, 140 mM NaCl, pH 7.40 at 37°C), a Pi stock solution (19.44 mM Na₂HPO₄, 4.56 mM NaH₂PO₄, 100 mM HEPES, 140 mM NaCl, pH 7.40 at 37°C), and a protein solution containing 1 mg/mL fetuin-A and 50 mg/mL albumin (Probumin, Millipore, #82-045-1) in saline.

Using a Liquidator 96™ pipetting system (Rainin, Mettler-Toledo) in a temperature-controlled room (34.5°C), the plate was prepared through a sequential addition and mixing process to ensure homogeneity. First, 20 µL of saline was dispensed per well, followed by 80 µL of protein solution containing different amounts of Mg and a one-minute shaking interval. Next, 50 µL of Pi stock was added with another minute of shaking, and finally, 50 µL of Ca stock was added, completing the formation of a supersaturated mineral solution, followed by a final one-minute shake. The final concentration was 140 mM NaCl, 50 mM Hepes 0.4 mg/mL fetuin-A, 20 mg/mL albumin, 10 mM Ca, 6 mM Pi, and 0 – 4 mM Mg.

Turbidity was continuously monitored using a Nephelostar® plate reader (BMG Labtech, Ortenberg Germany), with its internal heater disabled to maintain an internal temperature of 36.5–37°C. The instrument was programmed to perform 200 measurement cycles. Each cycle lasted 180 seconds and included a 1.5-second reading per well with a laser intensity of 85%, gain 85%, and a beam focus of 1.5 mm. The resulting relative nephelometric unit (RNU) values for each well were transferred to Microsoft Excel for preparation, and GraphPad Prism where nonlinear regression analysis log(agonist) vs. response–variable slope with four parameters (robust fit method) was applied to the curves to derive the T50 time and RNU values.

### Calcium phosphate precipitation inhibition (CAPPI) assay

The CAPPI assay was developed to quantify how Mg influences the stability of Ca and Pi in protein-rich solution in preventing sedimentation by centrifugation. The assay measures the ability of complex solutions like serum-containing cell culture media, which contain the mineral chaperone fetuin-A and BSA, two major components of fetal bovine serum (FBS), to form stable protein-mineral complexes that do not pellet during 10 min centrifugation at 20,000 x g. Magnesium stock solution (1M in water) was added to DMEM containing 10% FBS to bring the final concentration to 0.8, 3, or 9 mM total Mg. Ca and Pi stock solutions (0.5M in water) were consecutively added with intermittent mixing to a total volume of 200 µL, resulting in a final concentration of 4.3 mM Ca and 3.0 mM Pi. We added mineral in the following order, with thorough mixing between each addition to avoid local supersaturation: base medium including protein (FBS), then Mg, then Pi, then Ca.

Samples were incubated at 37°C for the times indicated in the figures and then centrifuged (10 min, 20,000 × g, 4°C) to separate the supernatant (soluble mineral and protein-mineral complexes) from the pellet (protein-mineral and mineral pellet). To quantify mineral in each fraction, 100 µL of supernatant was mixed sequentially with 50 µL of 0.6 M HCl and 50 µL of ammonium buffer (45 mM NH₄Cl pH 10.5, 5% v/v NH₄OH) to dissolve mineral and neutralize pH, respectively. The remaining 100 µL (containing the pellet) was dissolved by adding the same amount of HCl for dissolution and NH₄Cl solution for pH neutralization. Mineral content (Ca, Pi, Mg) in both fractions was determined by routine diagnostics at RWTH Aachen University Clinics Diagnostics Center (LDZ).

### Particle characterization by nanotracking analysis (NTA)

Particle size and concentration were analyzed using a NanoSight NS300 system (Malvern Panalytical) equipped with a 405 nm laser. Samples were diluted 1:250 to 1:1,000 in a buffer (140 mM NaCl, 20 mM HEPES, 1.8 mM CaCl₂, 0.9 mM Pi, 0.8 mM MgCl₂, pH 8.0) to achieve optimal particle concentration for analysis. For each sample, five 1-minute videos were recorded (n=5) and analyzed to determine the size distribution and concentration of protein-mineral complexes known as calciprotein particles (CPP). A 1 ml flow cell and a Blue488 laser were used. Camera level was set to 12, detection threshold was set to 5.

### Vascular smooth muscle cell culture

Immortalized human vascular smooth muscle cells, VSMC [20] clone IM3 were seeded into 24-well plates at a density of 10,000 cells per well and cultured for 7 days in DMEM (Gibco, 31053-044, 1.8 mM Ca, 0.9 mM Pi, 0.8 mM Mg) supplemented with 10% FBS (Pan Biotech, P30-3033), 1% Penicillin-Streptomycin (Gibco, 15140-122), and varying concentrations of additional Ca, Pi, and Mg.

### EBM-based calcification medium

An endothelial calcification medium (ECM-2) was formulated based on the commercial endothelial base medium EBM-2 (Lonza) by adjusting to 10% FBS, 4.3 mM total Ca, and 3.0 mM total Pi as in calcification medium (CM). We never observed any cell calcification in ECM-2. The EBM-2 base medium was analyzed for mineral concentrations, and 9 mM total Mg was detected. Therefore, we tested the influence of 0-10 mM added Mg to DMEM-based CM for calcification of VSMCs.

### DMEM-based calcification media

Our cell-based calcification studies employed DMEM (10% FBS) with varying additional Ca, Pi, and Mg. We varied mineral supersaturation over a wide range to test the effect of Mg on the stability of DMEM-based cell culture media highly supersaturated in Ca and Pi. SFigure 1 shows the full factorial design with Mg concentrations of 0, 1.25, 2.5, 5.0, or 10 mM added to a wide range of combinations of Ca and Pi. We added mineral in the following order, with thorough mixing between each addition to avoid local supersaturation: base medium including protein (FBS), then Mg, then Pi, then Ca. Control wells with no additional Ca or Pi were included to assess the effect of additional Mg alone.

**SFigure 1.**
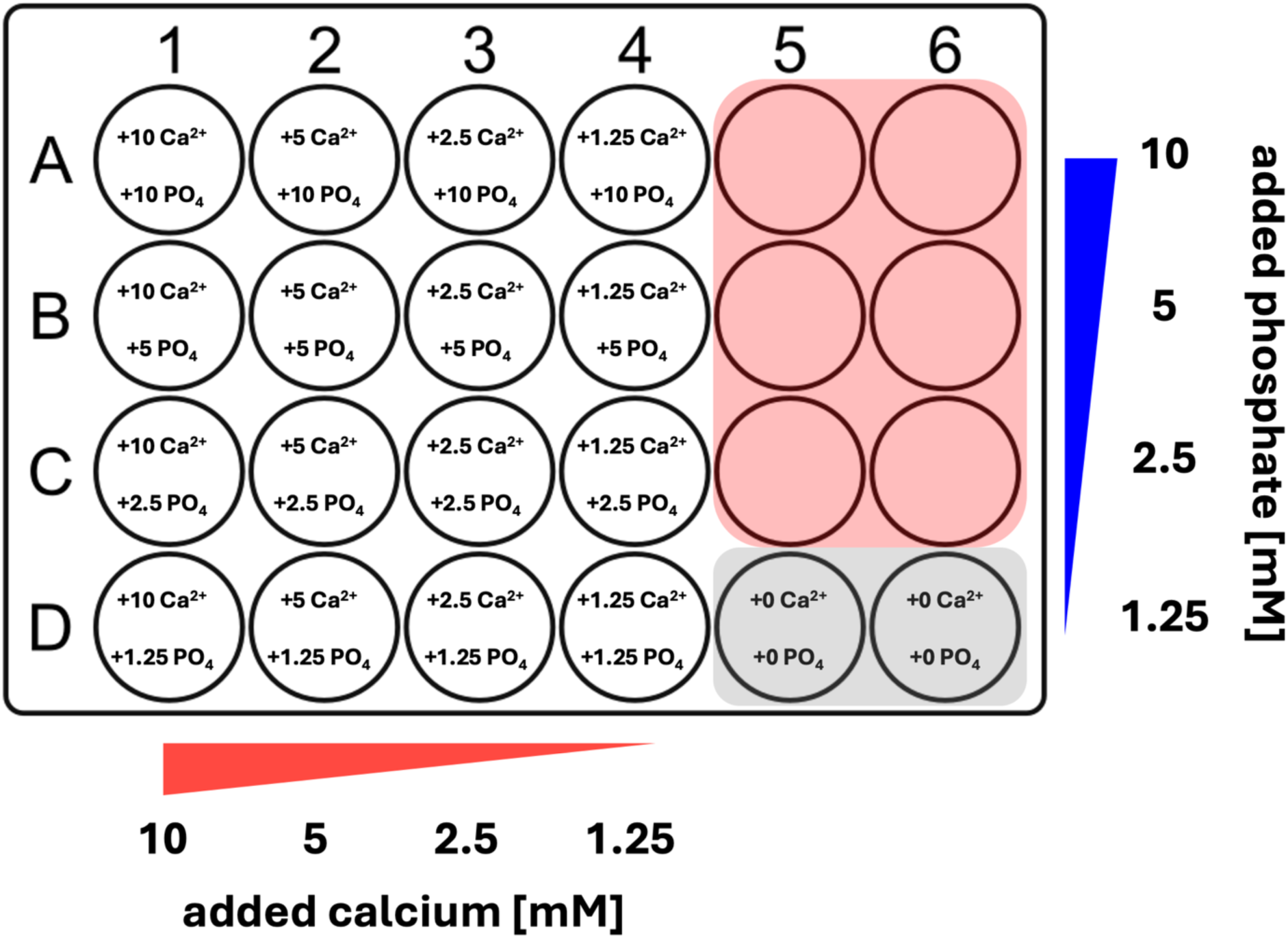
Factorial design for calcium, phosphate, and magnesium exposure in cell culture. Calcium and phosphate are added to DMEM/10% FBS in each well at the indicated amounts. Replica 24-multi-well plates were prepared with identical added Mg for each well. Added Mg concentrations tested were 0, 1.25, 2.5, 5, and 10 mM Mg resulting in five 24-multi-well plates per complete experiment. Wells in the red area were not used. Wells in the gray area had no added Ca and Pi, containing only 0-10 mM additional Mg.

### Live imaging and quantification of VSMC calcification with fluorescence-labelled fetuin-A

For live, longitudinal imaging of calcification, recombinant fluorescent fetuin-A-mRuby (100 µg/mL) was added to cell culture media as described [18]. A live imaging system (Cellcyte X, Cytena) was used to monitor calcification in real-time. Cells were imaged every 3 hours using brightfield and red fluorescence channels (exposure: 2000 ms, gain: 8 dB). Four fields of view were recorded per well using a 10x objective. The full-frame fluorescence intensity, proportional to fetuin-A-bound mineral deposition, was quantified using Cellcyte Studio software (Cytena) and exported in .csv file format for further analysis.

### Calcification and cell viability staining

For end-point observations, cells were incubated at the end of their culture period with recombinant fluorescent fetuin-A-mRuby (100 µg/mL) for 1 hour at 37°C to label mineral deposits as described [18]. Wells were then washed with serum-free DMEM and counter-stained with 50 µg/mL fluorescein diacetate (FDA, Sigma-Aldrich, 596-09-8) in serum-free DMEM for 30 seconds to label live cells. FDA staining solution was replaced with serum-free DMEM, and cells were immediately imaged using a Leica DMI 6000 inverted microscope and a 10x objective. Alternatively, C.Live Tox green stain (Cytena) was used to stain nuclei of dead cells without harming live cells.

### Alizarin Red staining

Cells were fixed with 4% paraformaldehyde for 30 minutes at room temperature and washed three times with PBS. Cells were then stained with 250 µL of 40 mM Alizarin Red S solution (Sigma-Aldrich, A5533; pH 4.2) for 10 minutes on a plate shaker. After staining, wells were washed with 1 mL distilled water until the washes ran clear. Cells were imaged again using a Leica DMI 6000 inverted microscope and a 10x objective. Four neighboring overlapping images were recorded and combined into a single image using Fiji software [21].

### Alizarin Red quantification through cetylpyridinium chloride extraction

Alizarin Red extraction was performed using cetylpyridinium. 100 mM cetylpyridinium chloride solution was prepared by dissolving 0.34 g of cetylpyridinium chloride (Sigma Aldrich, cat. C0732) in 10 mM sodium phosphate buffer. Sodium phosphate buffer was prepared by combining 81% disodium hydrogen phosphate and 19% sodium phosphate monobasic in distilled water. The solution was mixed using a magnetic stirrer until completely dissolved. The pH was then adjusted to 4.2 using 0.6 M HCl. The solution was freshly prepared and used within 24 hours.

Once prepared, 1 mL of 100 mM cetylpyridinium solution was added to each well of a 24-well plate. Plates were incubated at 37°C for 30 minutes. Multiwell plates were placed on a plate shaker to agitate and mix the solution. 100 µL of solution were taken from each well and transferred to a transparent 96-well plate for plate reader measurements. The absorbance was measured at 560 nm against a phosphate buffer blank.

### Simulated effect of magnesium on Alizarin Red staining intensity

The A560 values obtained with cetylpyridinium extracts were imported into MATLAB (The Mathworks Inc.) for interpolation. All measured conditions were assembled into a matrix alongside their measured extract readings. For all tested Mg concentrations, a third-degree polynomial fit was used to calculate Alizarin Red response surfaces. Each calculated surface consisted of 1000 data points. The quality of the fit of each surface was assessed and after each surface was found to be sufficiently fit (R^2^ > 0.95), the surfaces were used to calculate a unified function that predicted an Alizarin Red staining intensity for any combination of Ca, Pi, and Mg concentrations in the range. The unified function was used to calculate the maximized surface for simulated Alizarin Red staining, indicating the Mg concentration that was predicted to yield the highest intensity Alizarin Red staining, and thus the most calcification, for a given calcium and phosphate concentration.

## Results

### Stability of saturated calcium phosphate solutions - Calcification propensity test (T50) shows that Mg delays CPP-1 to CPP-2 phase transition

First, we performed T50 calcification propensity tests for samples comprised of buffer containing serum proteins fetuin-A (0.4 mg/mL) and albumin (20 mg/mL), with 10 mM Ca, 6 mM Pi, and increasing concentration of Mg (0 – 4 mM) measured over 10 hours. The T50 time indicates the time required for the sample to reach 50% of the total increase in turbidity caused by calciprotein particle CPP phase transition through Ostwald ripening from spherical particles with amorphous mineral called primary CPP-1 to elongated particles with crystalline mineral called secondary CPP-2. At the beginning of the measurement, sample turbidity was measured at roughly 150-200 relative nephelometric units RNU (Figure 1). After completing the phase transition, all samples measured between 300 and 600 RNU. Once the phase transition was complete, the samples maintained a constant turbidity value as the transition had formed stable protein-mineral CPP-2 phases. For a visual representation we performed the T50 assay in 1 mL cuvettes instead of microtiter plates. SMovie 1 shows that the formation of colloidal protein-mineral complexes was instantaneous. This prevented precipitates from falling out of solution, which happened in the protein-free incubation. Hardly any difference was visually apparent in the final turbidity of CM with increasing Mg in the cuvette-based assay shown in SMovie 1. In contrast, Mg dose-dependently stabilized CPP-1 in the microtiter plate T50 assay shown in Figure 1, thereby delaying the transition of amorphous CPP-1 to crystalline CPP-2. At the lowest Mg concentration (0.8 mM total), the T50 time was around 75 minutes. The T50 time was increased by increasing Mg concentration with 0.5, 1, 2, 3, and 4 mM added Mg to approximately 100, 125, 200, 300, and 450 minutes, respectively.

**Figure 1.**
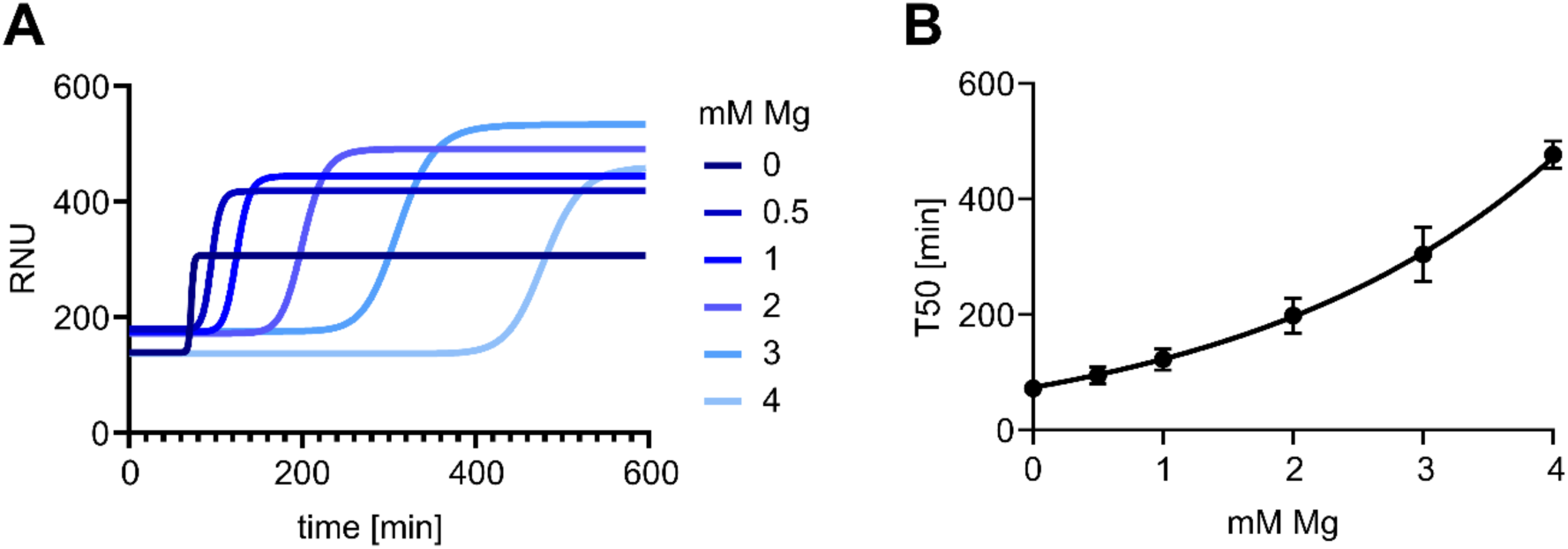
Stability of CPP protein-mineral complexes in solution measured by T50 assays. A buffered solution containing 140 mM NaCl, 50 mM Hepes purified fetuin-A (0.4 mg/ml), bovine serum albumin (20 mg/mL), 10 mM Ca, 6 mM Pi, and Mg from 0 – 4 mM was prepared. (A) Turbidity was continuously recorded for 10 hours using a plate nephelometer and the T50 times marking the transition from CPP-1 to CPP-2 were determined by non-linear curve fitting. (B) T50 times of each Mg concentration condition, showing a clear increase in T50 time with increasing Mg concentration. RNU, relative nephelometric units. Each measurement was performed in triplicate (n = 3). Error bars depict standard deviation with very small errors not visible.

**SMovie 1.**
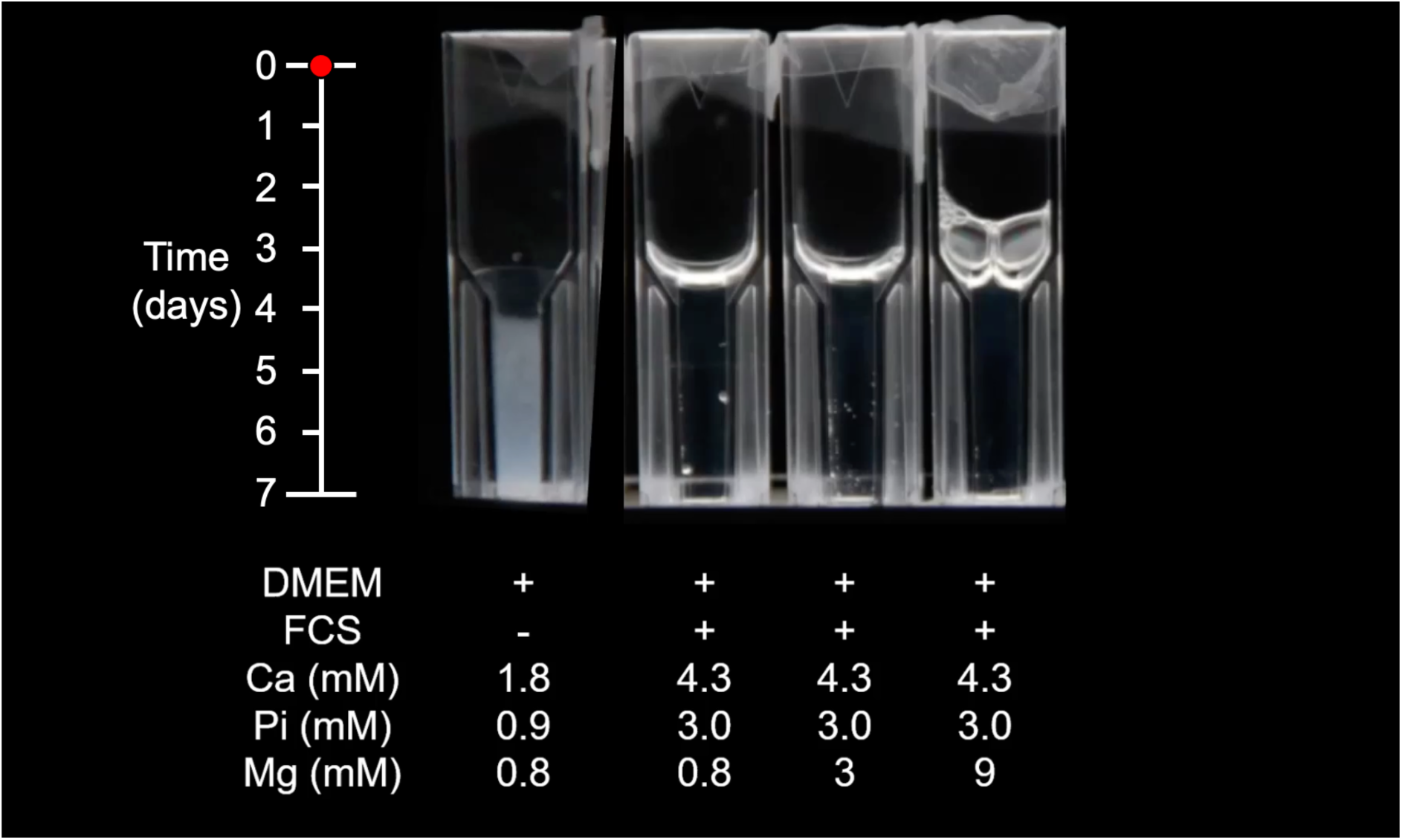
Effects of serum proteins and magnesium concentration on stability of calcium-phosphate particles in DMEM. DMEM with 4.3 mM Ca, 3.0 mM Pi, and 0.8, 3, or 9 mM Mg was mixed in couvettes and incubated at 37C° for 8 days (196 hours). One sample contained DMEM without FBS (left) while the other samples contained DMEM+10%FBS with 0.8, 3, or 9 mM Mg. Images were acquired every 3 hours and combined into a movie running at 10 frames per s. In DMEM without FBS, immediate precipitation occurred within the first hour. In DMEM with FBS, regardless of magnesium concentration, stable colloidal particles were formed and remained in solution for the duration.

### Calcium-phosphate precipitation inhibition (CAPPI) assay shows a lack of magnesium co-precipitation

Next we employed calcium phosphate precipitation inhibition (CAPPI) assays, measuring the mineral content of the supernatant and centrifuged pellet formed in DMEM with 10% FBS, 4.3 mM Ca, 3.0 mM Pi, and 0.8, 3, or 9 mM total Mg. Shown are the Ca (Figure 2A-C), Pi (Figure 2D-F), and Mg (Figure 2G-I) content of the supernatant (blue) and centrifuged pellet (red). All assays contained sufficient protein to increase equilibrium Ca and Pi at the end of the measurement period from 1 mM Ca, 0.5 mM Pi in protein-free CAPPI mixtures (not shown) to 2.5 mM Ca and 1.5 mM Pi regardless of Mg concentration. Thus, CAPPI complements the T50 assay (Figure 1) which showed that Mg strongly delayed the transformation of CPP-1 to CPP-2, both of which are suspended as colloids. CAPPI allows an estimate of non-pelletable low density protein-mineral complexes that are invisible in the T50 assay. At the 4.3 mM Ca and 3 mM Pi initial supersaturation of this particular CAPPI setup, roughly half of the Ca and Pi obviously existed as ionic or very low density protein-mineral complexes that could not be pelleted even by 10 min centrifugation at 20,000 x g, while the other half of the mineral formed higher density complexes which pelleted.

**Figure 2.**
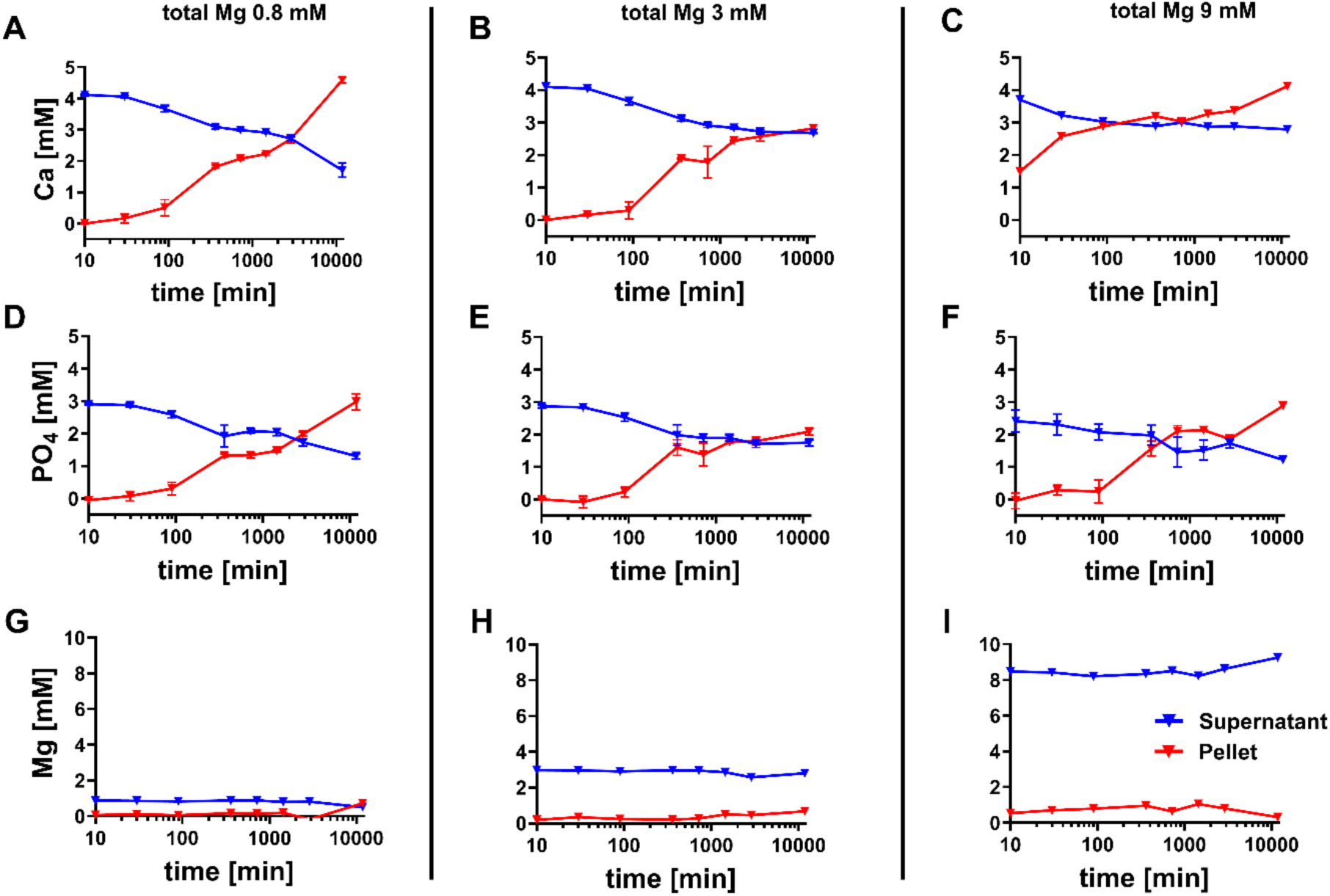
Stability of protein-mineral complexes in solution measured by calcium phosphate precipitation inhibition (CAPPI) assays. DMEM calcification cell culture medium with 10% FBS, 4.3 mM Ca, 3.0 mM Pi, and varying total Mg concentration (A,D,G: 0.8 mM Mg; B,E,H: 3 mM Mg; C,F,I: 9 mM Mg) were incubated at 37 °C, 5% CO_2_ for the indicated time. High density mineral complexes were pelleted by centrifugation at 20,000 x g, 4 °C for 10 min. Mineral contained in pellets is shown in red, while mineral in the supernatant is shown in blue. (A-C) Calcium content of pellet and supernatant. (D-F) Phosphate content of pellet and supernatant. (G-I) Magnesium content of pellet and supernatant. Each measurement was performed in triplicate (n = 3). Error bars depict standard deviation with very small errors not visible.

With time, Ca and Pi in the supernatant decreased, and Ca and Pi in the pellet increased in each mixture suggesting slow and continuous formation of pelletable protein-mineral complexes from low density precursors. Increasing total Mg in the CAPPI mixture did not change the kinetics and amount of pelletable mineral. Mg did not dose-dependently increase in the pellet like Ca and Pi did, even at 9 mM total Mg in the assay, suggesting that Mg did not overtly participate in the formation of pelletable mineral and protein-mineral complexes.

### Nanoparticle Tracking Analysis (NTA) shows that Mg influences protein-mineral particle size and number

Figure 3 shows the mean and mode for hydrodynamic diameter, as well as the concentration of protein-mineral particles/mL. Mean and mode diameter of particles formed in the presence of increasing Mg content were similar at earlier timepoints but diverged by 170 hours / 7 days. Figures 3A,B show that the mean and mode of the particle diameter in medium without added Mg (0.8 mM) slightly increased over the first 48 h and reached 150 nm (mean) and 125 nm (mode) respectively after 7 days. Particles derived from medium with elevated Mg (3 and 9 mM) had smaller mean and mode diameter throughout the experiment that never exceeded 100 nm diameter suggesting that elevated Mg prevented particle growth over one week. Particle concentration was also affected by additional Mg. Particle concentrations followed a bell-shaped kinetic indicating transient particle number increase and subsequent decrease. The particle concentration peaked around 0.5 hours (0.8 mM Mg) and 6-12 hours, respectively (3 and 9 mM Mg). The difference in particle concentration was most distinct at later timepoints. Figure 3C shows that by 7 days, medium containing 3 and 9 mM Mg, respectively had 2-3-fold more particles (8.8e10 ± 4.4e9 and 9.1e10 ± 6.0e9, respectively) than medium without additional Mg (3.3e10 ± 2.59e9). Collectively, these data suggest that protein-mineral particles initially grew by apposition until they transform from smaller colloidal particles into bigger solid particles that precipitate and escape measurement by NTA. Added Mg retarded both particle growth (Figure 3A,B) and reduction of suspended particle concentration (Figure 3C) providing smaller particles in greater numbers over 7 days.

**Figure 3.**
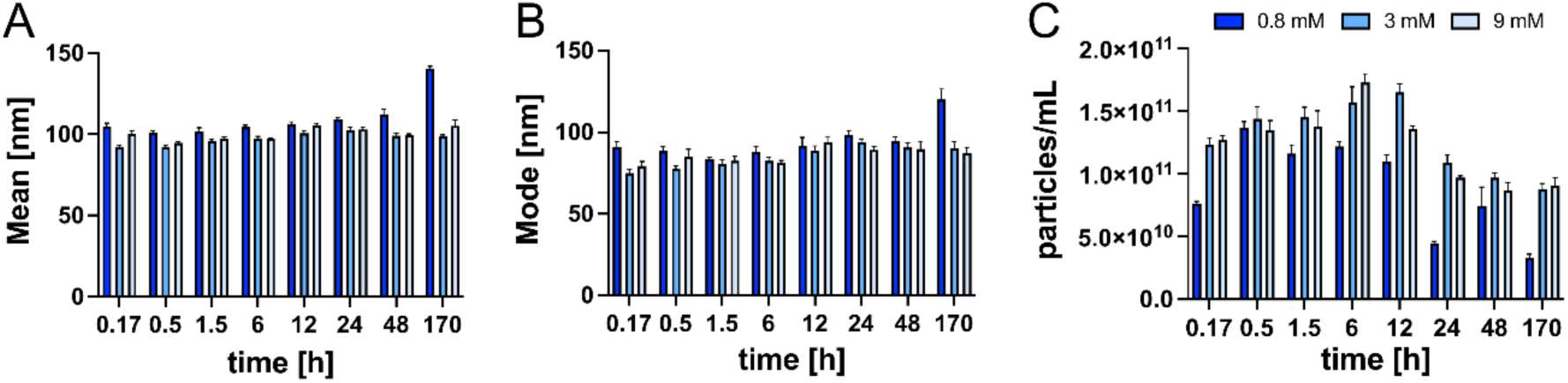
Nanoparticle tracking analysis (NTA) of protein-mineral particle size and number across varying magnesium concentrations. NTA results showing changes in particle hydrodynamic diameter mean (A), mode (B), and concentration (C) over 170 hours (7 days) in DMEM with 10% FBS, 4.3 mM total Ca, 3.0 mM total Pi, and 0.8, 3, or 9 mM total Mg (indicated by figure legend). Number of particles decreased over time due to particle aggregation and precipitation from the supernatant. Increased magnesium inhibited the increase in mean and mode diameter seen at 7 days. At all timepoints, 3 and 9 mM Mg had a higher concentration of particles and showed a significantly slower decrease in number of particles over time, indicating particle stabilization with slower particle aggregation and less precipitation. Bars represent mean±SD (n=5).

### Cell death in calcification media

Using Cellcyte X live cell imaging, continuous, longitudinal analysis of calcification in various calcification media employing live staining with high temporal resolution was made possible. We were able to elucidate differences between conditions containing varying amounts of Ca, Pi, and Mg. SMovie 2 shows IM3 vascular smooth muscle cells growing for up to 7 days without medium exchange in DMEM, DMEM-based calcification medium CM (DMEM, adjusted to 4.3 mM Ca and 3.0 mM Pi), or endothelial calcification medium EBM-2-based calcification medium ECM-2 (EBM-2, Lonza, adjusted to 4.3 mM total Ca and 3.0 mM total Pi). C.Live Tox Green staining was used to label the nuclei of dead cells. Due to the high proliferation rate of IM3 cells, a low initial cell density was used to avoid cell overgrowth and premature medium depletion during the 7-day culture period. SMovie 2 and the corresponding dead-cell quantification depicted in SMovie 2D show that cells in DMEM grew to confluency within 48 h. Around the same time cells grown in CM (SMovie 2B) showed cell death and associated dystrophic calcification of cell debris visible as opaque lesions in brightfield microscopy exacerbating between 42-72 hours. In contrast, cells grown in ECM-2 (SMovie 2C) did not calcify even after 7 days even though the medium contained the same amount of Ca and Pi like CM. The lack of observed calcification in cells cultured in ECM-2 is explained by the fact that EBM-2 basal medium contained approximately 9 mM Mg, which is sufficient to completely inhibit calcification. After 6 days, increasing cell death was observed in all conditions, which was due to medium nutrient depletion.

**SMovie 2.**
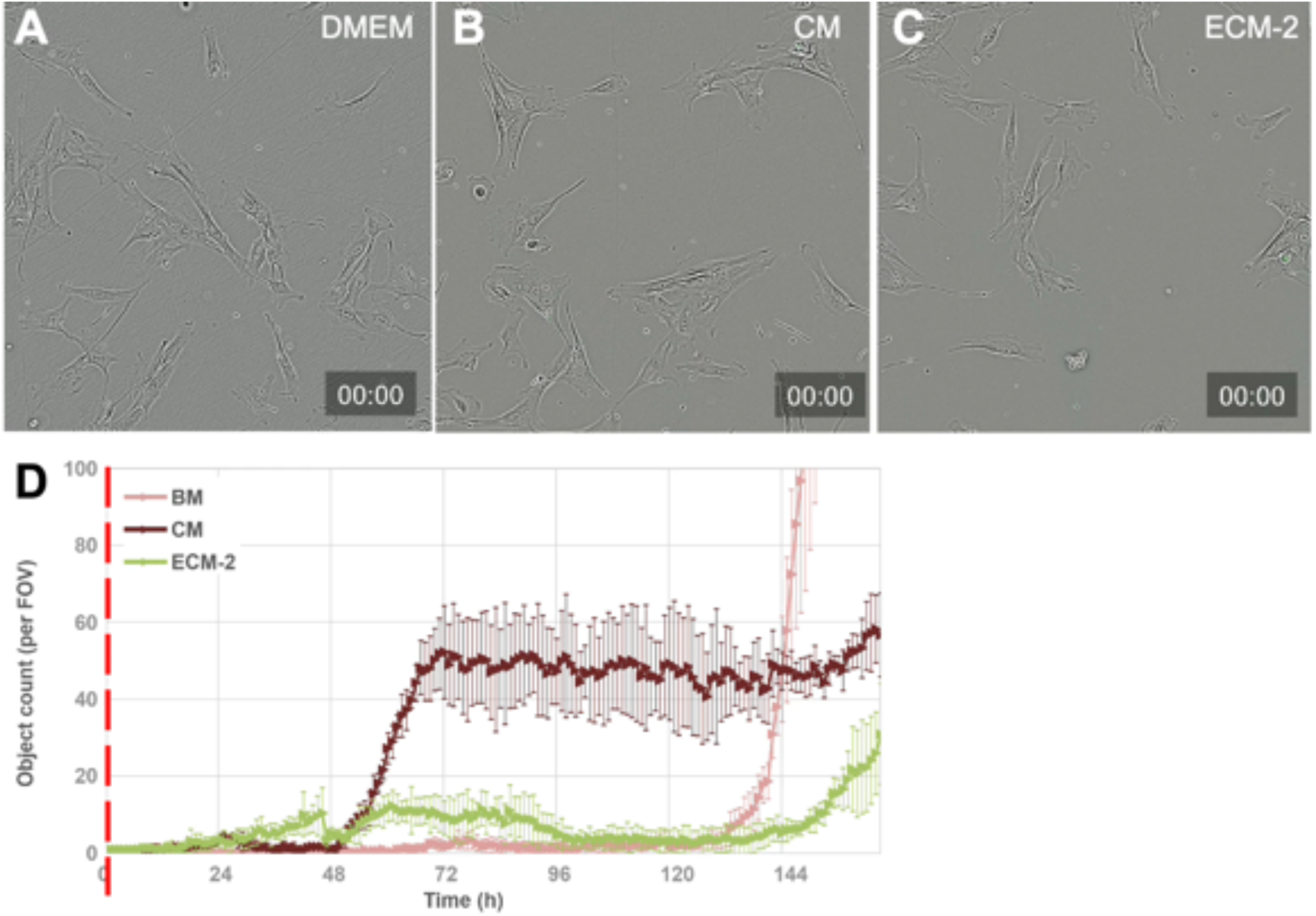
Live imaging movies showing cell death in various media. Cellcyte X live imaging of IM3 immortalized vascular smooth muscles over 7 days in (A) DMEM, 10% FBS, (B) calcification medium CM (DMEM, 10% FBS, adjusted to 4.3 mM Ca and 3 mM Pi), and (C) endothelial calcification medium ECM-2 (endothelial base medium EBM-2, 10% FBS, adjusted to 4.3 mM total Ca and 3.0 mM total Pi). Movies are time-lapse with frames acquired every three hours. C.Live Tox green, a nuclear stain demarcating dead cells, is visible in green. (D) Quantification of threshold object count in the green channel, indicating the number of dead cells per field of view. Each data series consists of four analyzed fields of view of the same condition. Each measurement was performed in quadruplicate (n = 4). Error bars depict standard deviation.

### Calcification Kinetics and Patterns are regulated by Mg and Pi

Using fluorescence-labelled fetuin-A, the development of calcification was also analyzed across conditions and timepoints. By acquiring one frame every 3 hours, sufficient temporal resolution was provided to determine differences in both the rate of development, and the morphology of the calcifications. SMovie 3 shows the developing calcification of smooth muscle cells over 7 days, indicated by fetuin-A-mRuby3 fluorescence. Magnesium delayed the onset of calcification in a dose-dependent manner. In addition, the morphology of the calcification changed with increasing Mg concentration. Lower concentrations of Mg (SMovie 3A) exhibited more rapid onset of calcification along cell boundaries, while higher concentrations (SMovie 3C) exhibited delayed onset followed by sudden and sheet-like calcification across the cell layer suggesting that calcification was predominantly driven by small protein-mineral particles that were more stable and therefore more abundant in media containing elevated Mg (Figure 3).

**SMovie 3.**
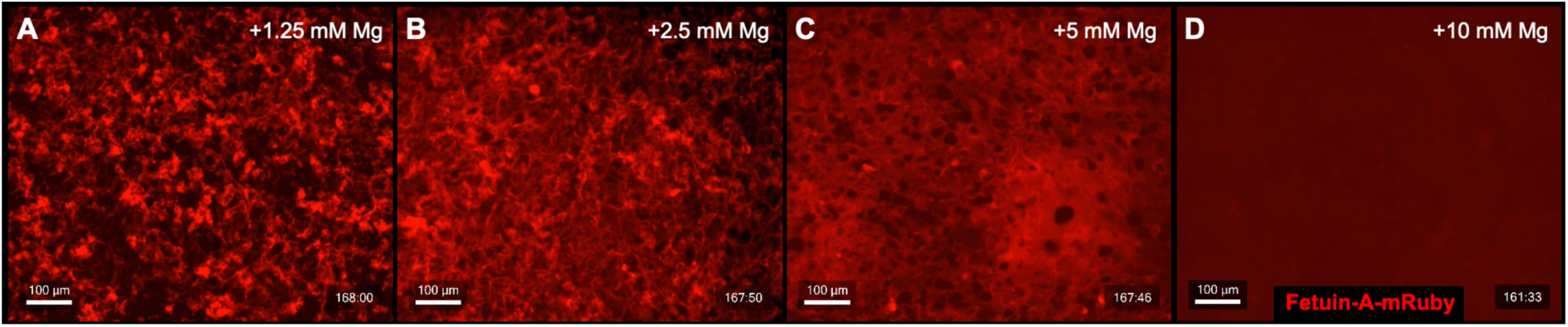
Live imaging movies showing calcification in varying magnesium. Cellcyte-X live imaging of IM3 immortalized vascular smooth muscle cells over 7 days in DMEM, 10% FBS, with 10 mM added Ca and Pi and (A) 1.25 mM, (B) 2.5 mM, (C) 5 mM, and 10 mM added Mg. Fetuin-A-mRuby staining of calcification is shown in red. Movement of living cells can be seen in SMovie 3A,B, but not in C. The morphology of the calcification can be seen to change with increasing Mg. Lower concentrations of Mg exhibited more rapid onset of calcification along cell boundaries, while higher concentrations exhibited delayed onset followed by sudden and sheet-like calcification across the cell layer. At the highest concentration, 10 mM added Mg, no calcification was observed.

Next, we systematically added Ca and Pi (+1.25-10mM) in DMEM/FBS medium with moderately increased Mg (+2.5 mM). In all high Ca conditions with low Pi FDA staining revealed confluent sheets of viable cells. In contrast, increasing added Pi from 1.25 mM to 10 mM in DMEM-based calcification media (0.9 mM Pi, 3.3 mM Mg, 11.8 mM Ca, all total) caused a marked decrease in viable cells after 7 days judged by FDA staining (SFigure 2A-C). The ECM was strongly calcified in all cases, yet morphology of calcification varied from predominantly cell boundary-associated at 1.25 mM additional Pi (2.15 mM total, SFigure 2D) to sheet-like calcification across the whole cell layer at 10 mM added Pi (SFigure 2D-F).

**SFigure 2.**
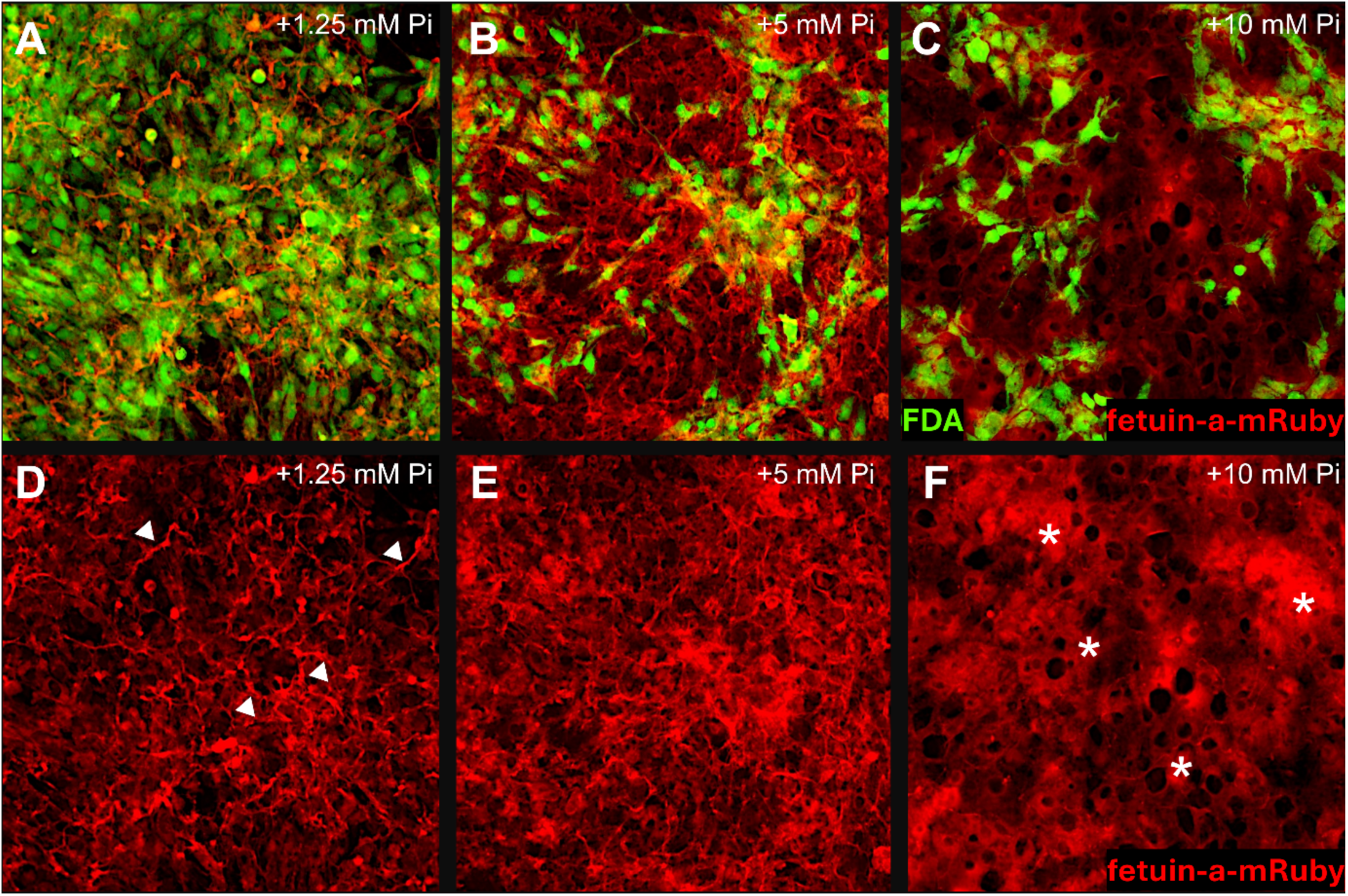
Fluorescence imaging showing calcification in varying phosphate. IM3 immortalized vascular smooth muscle cells over 7 days in DMEM, 10% FBS, with 2.5 mM additional Mg, 10 mM additional Ca, and 1.25 mM (A, D), 5 mM (B, E), and 10 mM (C, F) additional Pi. (A-C): FDA staining of viable cells is shown in green and (A-F). Fetuin-A-mRuby staining of calcification is shown in red. Lower concentrations of Pi exhibited calcification along cell boundaries (white arrows), while higher concentrations exhibited sheet-like calcification across the whole cell layer (white asterisks).

### Live imaging of calcification in cell culture using fluorescent fetuin-A-mRuby

Figure 4 illustrates the fluorescence intensity recorded during live imaging for calcification in two condition sets. Figure 4A shows conditions with 10 mM additional Ca and Pi across varying Mg concentrations, with Figure 4B showing a zoomed fit of the first 72 hours. Figure 4C shows conditions with 10 mM additional Ca and 5 mM additional Pi across various Mg concentrations, with Figure 4D showing a zoomed fit of the first 72 hours. In both conditions, the onset of calcification was delayed in a Mg dose-dependent manner. Ten mM added Mg completely prevented calcification in live cell culture even in the presence of 10 mM additional Ca and Pi, which would otherwise immediately precipitate onto cells. It was also observed that the higher Pi content in Figure 4A,B promoted faster calcification onset than the conditions containing less Pi (Figure 4C,D). In these lower Pi conditions, the progression of calcification was delayed such that the calcification took place over an extended period, while ultimately arriving at the same fluorescence intensity except in the highest Mg concentrations.

**Figure 4.**
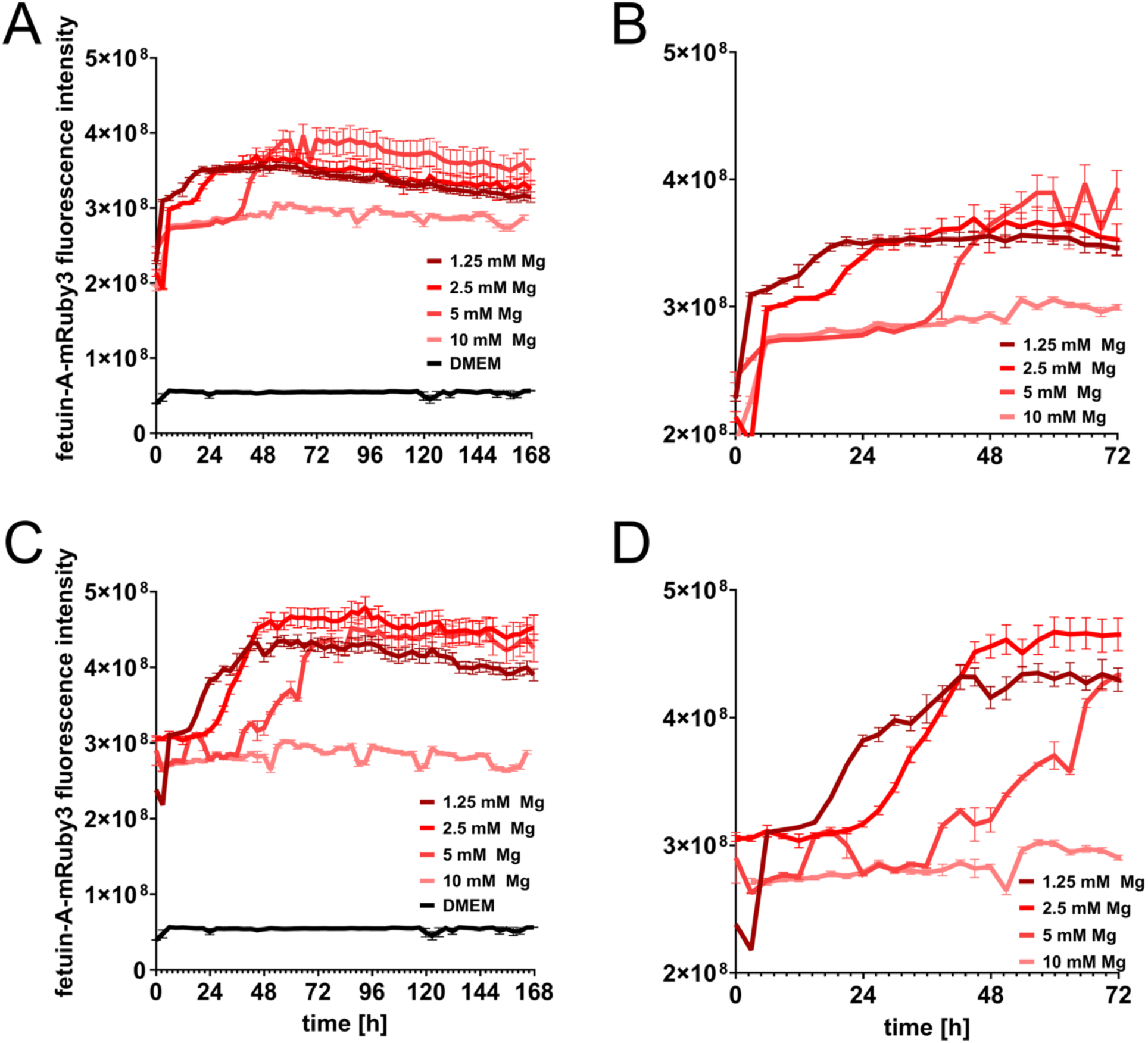
Live quantification of smooth muscle cell calcification in culture. Cellcyte X live fluorescence quantification of IM3 immortalized vascular smooth muscle cells over 7 days using fluorescence-labelled fetuin-A-mRuby3. Cells were cultured in DMEM, 10% FBS, with 10 mM additional Ca and either 10 mM (A-B) or 5 mM additional Pi (C-D) with additional Mg as listed in the figure legend. (A, C): Complete time course of live imaging data. (B, D): First 72 hours of time course, zoomed for readability. Each data series consists of fluorescence quantification of four fields of view per condition with error bars shown (n = 4). Fluorescence intensity of fetuin-A-mRuby3 indicates degree of calcification, with background intensity measured in DMEM. Error bars depict standard deviation with very small errors not visible.

### Calcification and cell viability staining

Figure 5 illustrates the endpoint analysis of calcifying vascular smooth muscle cells, showing calcification determined by fetuin-A staining and Alizarin Red staining (Figure 5A-O). After 7 days in calcifying conditions, the cells were stained with FDA (Figure 5P-T) and fetuin-A-mRuby (Figure 5K-O) to assess cell viability and the extent of calcification present in each condition. FDA staining indicated the presence of viable cells in all conditions, including those with the highest Ca and Pi content. Conditions with high phosphate had the least FDA-positive cells confirming phosphate toxicity (Figure 5P-T, top row). Low phosphate conditions also caused an apparent loss of FDA-positive cells. In these conditions however, the cells were lost because they proliferated quickly until the medium was exhausted and the cells began to contract and detach from the multi-well plate, causing loss of entire cell sheets (Figure 5P-T, bottom rows). All cells remaining attached nevertheless were fully viable and strongly FDA-positive. Fetuin-A-mRuby staining revealed nascent calcifications across all conditions, with more calcification in conditions with higher Ca. Alizarin Red staining (Figure 5F-J) showed that 1.25 mM – 5 mM additional Mg caused an increase in calcification especially at 10 mM Ca (Figure 5I). When additional Mg was increased to 10 mM (Figure 5J) calcification was inhibited at all conditions including 10 mM Ca and 10 mM Pi. The quantified extraction of Alizarin Red using cetylpyridinium (Figure 5A-E) corroborated the visual impression given by microscopy (Figure 5F-J). There was a Mg dose-dependent increase in the amount of extracted Alizarin Red at up to 5 mM added Mg, yet calcification was completely inhibited at 10 mM added Mg.

**Figure 5.**
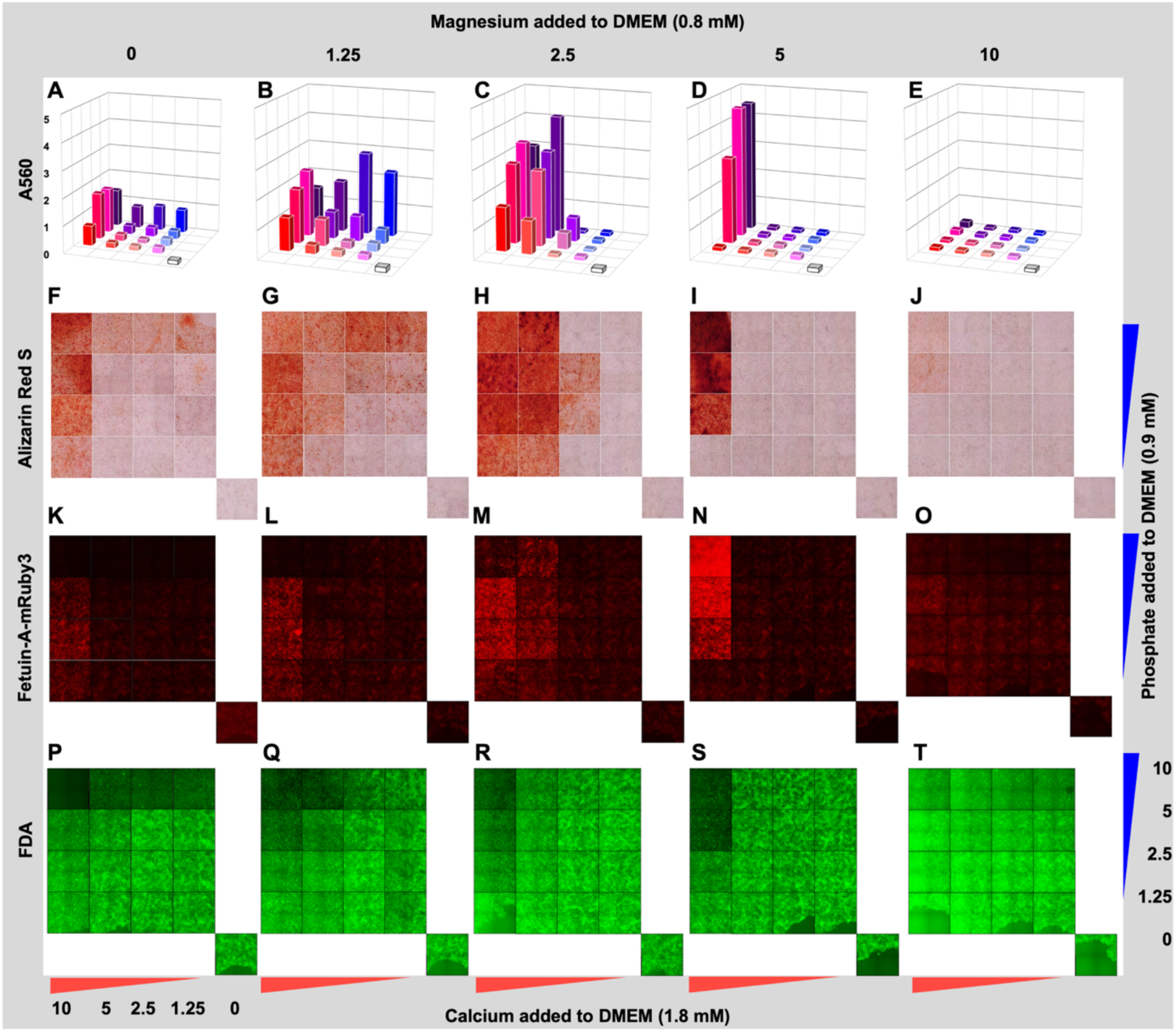
Endpoint analysis of cell viability, calcification, and Alizarin Red staining/quantification. Quantification and endpoint microscopy of IM3 immortalized vascular smooth muscle cells after 7 days in medium containing systematically varied amounts of Ca, Pi, and Mg. (A-E): 560 nm absorbance plate reader measurements from cetylpyridinium-extracted Alizarin Red for each condition. (F-J): Alizarin Red staining. (K-O): red fluorescent fetuin-A-mRuby3. (P-T): Fluorescein diacetate (FDA).

### Simulated effect of magnesium on Alizarin Red staining intensity

Figure 6 describes the relationship between Ca, Pi, and Mg concentration and the quantified Alizarin Red staining intensity shown in Figure 5. In Figure 6A, for each tested Mg concentration (indicated on the vertical axis) the predicted Alizarin Red staining intensity (indicated as red color) was calculated for all Ca and Pi concentrations in the range. The results show conditions where increasing Mg concentration produces more calcification than identical Ca and Pi concentrations at lower Mg concentrations. However, as Mg concentration continues to increase, a decrease in the predicted Alizarin Red value is observed until no calcification is predicted at 10 mM additional Mg (Figure 6A, top plane). Figure 6B shows a maximized, unified surface that predicts the Mg concentration (indicated by green color) that should provide the highest intensity Alizarin Red staining for any given Ca and Pi concentration within the range. By combining the interpolated planes into a unified function, the predicted Alizarin Red value for any given combination of Ca and Pi can be calculated. Maximizing this function gives the Mg concentration that is predicted to produce the highest Alizarin Red signal. Using this model, a total Mg concentration of approximately 4.5 mM was predicted to maximize the Alizarin Red value, and this calculated Mg concentration for maximum Alizarin Red staining, i.e. calcification, fell within a narrow range for all Ca and Pi concentrations, despite large variation in the simulated Alizarin Red staining intensity ranging from 2-9 absorption units.

**Figure 6.**
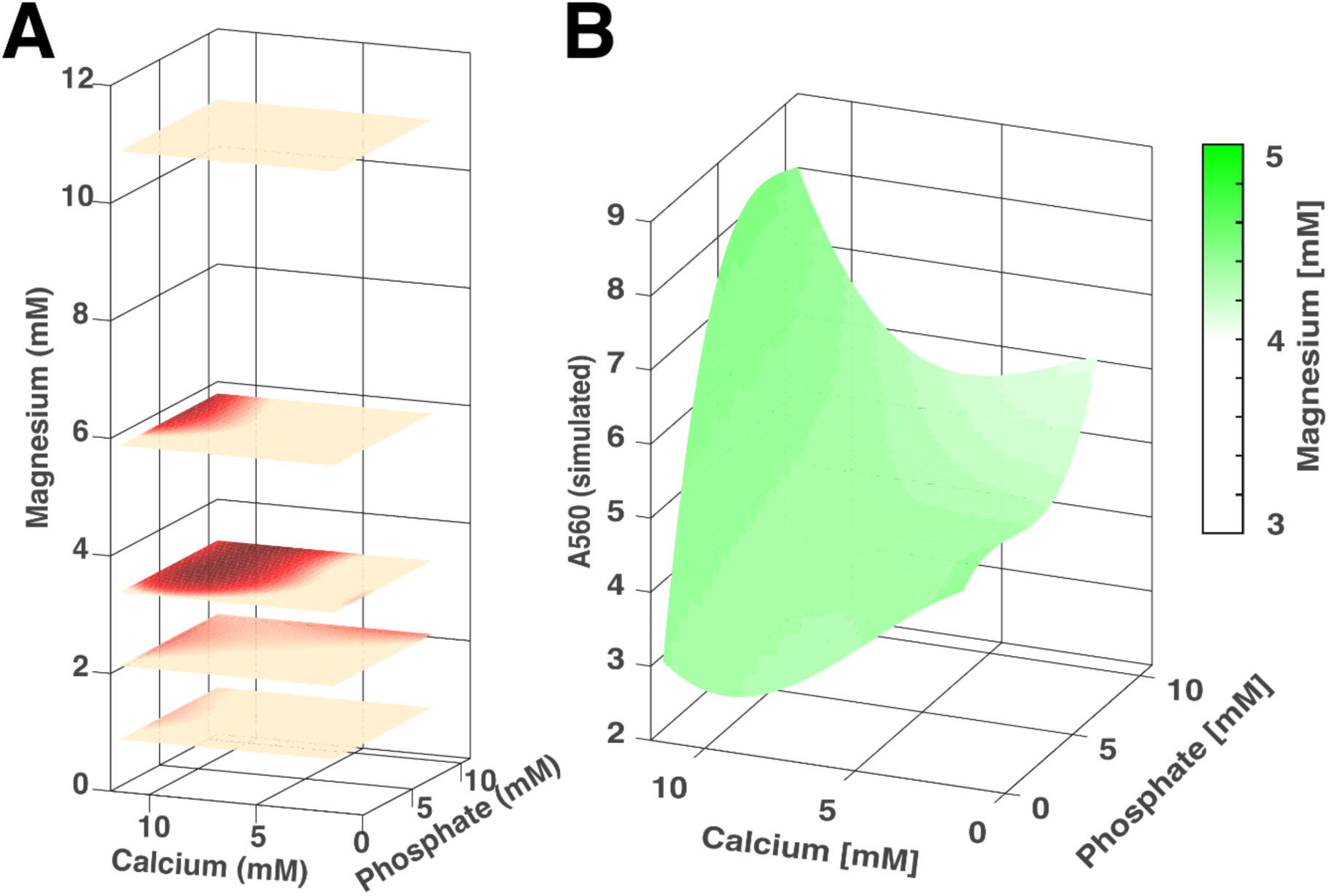
Simulated effect of magnesium on Alizarin Red staining intensity. (A), Seventeen 560 nm absorbance plate reader measurements from cetylpyridinium-extracted Alizarin Red were interpolated to 1000 values per Mg concentration using MATLAB and a third-degree polynomial fit. An increase in the predicted Alizarin Red value can be seen as the Mg concentration increases to 2.5 mM additional Mg (A, 3^rd^ plane from the bottom). (B), Maximized surface of the predicted Alizarin Red value for all conditions with matching Mg concentration indicated by green color. Approximately 4.5 mM was predicted to produce the most Alizarin Red staining at all Ca and Pi concentrations in the tested range.

## Discussion

Our work shows that Mg plays a dual role in ECM calcification, which is determined by its concentration, the degree of mineral supersaturation, and the presence of mineral-complexing proteins. Under the highly supersaturated yet stabilized conditions of our experiments, Mg’s function was dose-dependent. At moderate concentrations (1.25-5 mM added), Mg acted as a facilitator of matrix mineralization in calcification media by stabilizing protein-mineral phases as colloids that readily transfer mineral to a mineralizable ECM, as previously observed in the calcification of pure collagen fibers [13]. This process mimics the synthetic mineralization of the matrix by a process called polymer-induced liquid-precursor (PILP) [22]. In contrast, at 10 mM, Mg stabilized the solutions beyond a point where mineral could be released and appended to a substrate, which explains why endothelial cell medium containing 9 mM total Mg completely prevented cell calcification. Importantly, these effects are entirely determined by the medium composition, with cells making only a minor contribution beyond providing a calcifiable ECM.

The mechanisms underlying this dual role were investigated through several assays. The delay in calcification onset, evidenced by an increase in T50 time from 75 minutes at 0 mM Mg to 300 minutes at 3 mM Mg, aligns with Mg’s known capacity to adsorb to nascent crystal surfaces and distort hydroxyapatite lattice formation [23]. The calcium phosphate precipitation inhibition (CAPPI) assay confirmed that Mg reduces mineral precipitation without significantly incorporating into the pelleted phase, thereby sustaining supersaturation. There was some discrepancy in the recovery rate of Ca and Mg due to these ions competitively interfering in concentration measurement assays. Due to this interference, in combination with slight variations in handling, the final mineral concentrations had approximately ±10% error. However, nanoparticle tracking analysis revealed that Mg, in the presence of the mineral chaperone fetuin-A, suppresses particle aggregation and maturation with strong statistical certainty (p = 0.0007 when comparing 0.8 mM Mg with 3 mM Mg at 170 hours). By stabilizing the mineral phase in early-stage CPM and tCPP, which is echoed by the delay of transformation of amorphous CPP-1 into late-stage, crystalline CPP-2, Mg together with fetuin-A and other serum proteins preserves mineral in a highly dynamic colloidal state readily available for matrix calcification.

The inverse relationship between particle number and size under different Mg conditions is supported by the fact that overall protein-mineral particle volume is constant, when converting the numbers times the measured hydrodynamic diameters of particles into spherical volumes. The calculated total particulate volumes were similar (CM0, 4.85e16 nm³/mL; CM3, 4.43e16 nm³/mL; CM9, 5.53e16 nm³/mL) despite significantly different particle concentrations seen with increased Mg concentration. We attribute the discrepancy between total calculated volumes to combined measurement errors. It could also be explained by an increase in particles that are too small or insufficiently refractive to be detected. In any case, the shift in particle size distribution reflected a fundamental change in kinetics of the colloidal mineral phase.

We hypothesize that the observed stabilization of mineralization precursors relies on two synergistic shielding mechanisms operating at different scales. At the ionic level, Mg exerts its effect through competitive surface adsorption onto nascent calcium phosphate clusters. This ionic shielding not only raises the energy required for ion addition and growth but also introduces lattice strain that preferentially stabilizes the amorphous phases over the more stable, yet insoluble crystalline hydroxyapatite [15]. Simultaneously, protein stabilizes protein-mineral complexes through topological shielding. This involves the structural encapsulation of an amorphous mineral core by proteins like fetuin-A, forming a steric barrier to crystal growth in calciprotein particles (CPP) [17]. This protein encapsulation increased the stability of amorphous calcium phosphate and suppressed particle aggregation. The aggregation and maturation of CPP are further retarded by magnesium, increasing the lifetime of early CPP [24], especially of CPM and tCPP [13], which best sustain matrix calcification even by low complexity versions of CPP containing mainly fetuin-A [25] and albumin [26]. Thus, the early form CPM and tCPP are sufficiently supersaturated yet unstable enough to act as an ideal mineral “source” for a calcifiable “sink” like collagen fibers or ECM. The protein shielding may also act as a diffusion barrier preventing spontaneous mineral precipitation from solution yet allowing mineral transfer to calcifiable ECM [27]. Our data suggests that Mg, in addition to ionic shielding, enhances the efficacy of this topological shielding by promoting the formation of a more stable yet highly dynamic protein-mineral interfaces. The combined effect of ionic and topological shielding prolongs the colloidal suspension stage, increasing the circulatory lifetime and tissue penetrance of these mineralization-competent precursors. This explains the paradoxical increase in long-term mineral deposition at intermediate Mg concentrations. By preventing rapid, uncontrolled precipitation, the shielding mechanisms ensure a sustained delivery of preformed mineral to the cellular microenvironment, yet do not stop mineral transfer altogether.

The colloidal stabilization of calcium phosphate fundamentally shifts mineral accessibility for vascular smooth muscle cells. Live imaging with fluorescent fetuin-A-mRuby confirmed that Mg delayed calcification onset in cell culture, yet endpoint analysis revealed an increase in Alizarin Red staining at intermediate Mg concentrations (2.5-5 mM added) under these specific mineral conditions. To further explore this paradoxical effect, we mathematically modelled the Alizarin Red staining response across the entire experimental range of Ca, Pi, and Mg concentrations. By generating third-degree polynomial fit response surfaces (R² > 0.95), we simulated the staining intensity for any condition within the tested range. Strikingly, this simulation revealed that the Mg concentration associated with maximum calcification signal consistently covered a narrow window around 4.5 mM, independent of the specific Ca and Pi concentrations. This finding underscores Mg’s dual role: we hypothesize that this narrow "sweet spot" represents the concentration at which the promotional effect of Mg on cellular activity is maximized, before being surpassed by the dominant role as a mineral stabilizer and potent calcification inhibitor at higher concentrations. Using this information, an ideal calcification medium can be formulated for any cell line using the method described in this work. According to the simulation, an ideal calcification medium based on inorganic compounds (no ß-glycerophosphate) would contain roughly 10-12 mM total Ca, 6-8 mM total Pi, and 4.5 mM total Mg. Further refinement of this formulation would be possible by exploring combinations near these predicted concentrations. Fluid preparations containing these minerals must be mixed carefully to avoid local supersaturation and immediate precipitation due to high mineral concentration in stock solutions. For serum-containing media, mineral should be added in the following order, with thorough mixing between each addition: base medium including protein (FBS), then Mg, then Pi, then Ca.

Combining live imaging of fluorescent fetuin-A with nanoparticle tracking and endpoint assessments captures calcification kinetics previously inaccessible through single-timepoint assays. The real-time delay in fluorescence onset combined with the colloidal dynamics resolved by NTA provides a cohesive model: Mg decouples mineral nucleation from cellular processing by enhancing the stability of CPP in solution. Clinically, these insights help contextualize the potential U-shaped epidemiological associations of serum Mg with cardiovascular risk [28]. Low Mg permits rapid pathological precipitation, moderate Mg enables cell-driven mineralization potentially contributing to chronic calcification burden, and high Mg imposes near-complete inhibition of calcification.

Given these findings, we propose that Mg-stabilized CPM and tCPP act as a sustained mineral reservoir for ECM calcification, either directly, or after endocytosis of late CPP forms (CPP-1 and CPP-2), lysosomal disassembly and reassembly of early labile CPM and tCPP upon exocytosis [13]. This highlights a critical balance, where low to moderate Mg levels (2.5-5 mM added) enhance mineral bioavailability—particularly in high-Ca environments—while high levels (>5 mM) suppress both passive precipitation and cell-driven mineralization. This aligns with clinical observations linking hypomagnesemia (<0.7 mM serum Mg) to accelerated vascular calcification in chronic kidney disease [29]. In contrast, increased dietary Mg was shown to rescue severe soft tissue calcification in mice [30, 31].

While these findings provide some clarity to the onset and progression of calcification in cell culture, simple cell culture of immortalized vascular smooth muscle cells cannot possibly replicate the behavior of primary cells or even complete tissues from chronic kidney disease patients, particularly regarding disease-specific pathways of osteogenic differentiation. Furthermore, the simplified DMEM/FBS system excludes the complex milieu of cellular clearance of circulating mineral by the kidney, liver and spleen [19, 32], uremic toxins, cytokines, and hormonal influences prevalent in chronic kidney disease that likely modulate cell calcification. Validating these findings in vivo using chronic kidney disease models with variable dietary Mg intake, while correlating serum T50 times with calciprotein particle size and distribution, is essential to confirm physiological relevance [33]. Exploring combination therapies employing Mg with established phytate or inositol-based calcification inhibitors, [34, 35] might further harness colloidal stabilization of protein-mineral complexes to enhance therapeutic efficacy.

Collectively, our findings render Mg a powerful experimental modulator of calcification kinetics in cell-based systems—not merely a passive inhibitor, but an active colloidal participant that controls the transition from soluble mineral to cellular mineralization. By stabilizing bioavailable amorphous protein-calcium phosphate complexes and CPP, Mg also creates an extended temporal window where protein-mineral complexes remain accessible to clearance while delaying precipitation. In vivo, this controlled stabilization buys time for clearance of early stage protein-mineral complexes CPM to prevent buildup of more stable and potentially harmful late-stage CPP [19]. This stabilization also improves in vitro calcification experiments: it decouples passive mineralization from cell-mediated processes, synchronizes calcification progression across cell populations, and generates well-defined kinetic phases amenable to temporal analysis. The prolonging effect enables precise time-targeted sampling for cell studies including advanced imaging or single-cell sequencing, allowing researchers to capture transient cellular states during calcification initiation and progression that would otherwise be obscured by rapid mineral precipitation. Collectively the results elevate Mg from a simple culture additive to a useful tool for resolving the mechanics of cell-mediated mineralization pathways.

## Supporting information

Smovies 1-3

## Acknowledgments

This work was funded by the Deutsche Forschungsgemeinschaft (DFG, German Research Foundation) – TRR 219 – Project-ID 322900939 (WJ-D and CG) and Project-ID 40304155 (WJ-D).

